# PanKbase Integrated Single-Cell Map: A Comprehensive Atlas of Human Pancreatic Islets

**DOI:** 10.64898/2026.06.02.729719

**Authors:** Ha T.H. Vu, Han Sun, Parul Kudtarkar, Seth A. Sharp, Liza Brusman, Fan Feng, Thomas Bate, Julie A. Jurgens, Yiqun Wang, Yuanhao Huang, Runbo Mao, Sierra Corban, Amanda K. Huber, Alex Shilin, Ying Sun, Sara Narayanaswamy, Dongkeun Jang, Catherine C. Robertson, Shristi Shrestha, Trang Nguyen, Patrick Smadbeck, Lu Zhang, Mackenzie Brandes, The PanKbase Consortium, Jason Flannick, Noel Burtt, Shuibing Chen, Jie Liu, Jean-Philippe Cartailler, Benjamin F. Voight, Michael L. Stitzel, Marcela Brissova, Anna L. Gloyn, Kyle J. Gaulton, Stephen C.J. Parker

## Abstract

Single-cell RNA sequencing (scRNA-seq) of human pancreatic islet tissue is a powerful tool for investigating type 1 diabetes (T1D). However, individual datasets are limited in size and fragmented across donors, laboratories, and experimental conditions. To address this, we constructed a comprehensive, integrated scRNA-seq atlas of isolated human pancreatic islets by collating publicly available data generated from tissue provided by resources including the Human Pancreas Analysis Program, the Integrated Islet Distribution Program, and Prodo Labs. Systematic quality controls were implemented to select high-quality samples, reads, and cells.

During integration, we accounted for important variables such as age, sex, body mass index, origin study, treatments, islet distribution resources, and sequencing chemistry. Our single-cell atlas comprises 191 high-quality samples from 140 donors (59 female, 81 male) across five phenotypic groups: no diabetes (controls, n=69), autoantibody positivity without diabetes (n=12), pre-diabetes (n=11), T1D (n=12), and type 2 diabetes (T2D) (n=36). In total, the atlas contains 448,935 cells, capturing 13 distinct populations, including alpha cells (43.3%) and beta cells (26.8%), as well as groups such as immune cells (0.6%). Publicly available at www.pankbase.org, this atlas provides a platform for hypothesis-driven investigation of diabetes pathophysiology and, given rigorous quality control, is well-suited for downstream machine-learning applications.

**Article Highlights:**

- Current scRNA-seq datasets of pancreatic islet tissue are limited in size and scattered across donors, laboratories, and experimental conditions, underscoring the need for a consolidated resource.
- We harmonized datasets from multiple sources to build a comprehensive single-cell map of isolated human pancreatic islets.
- Our atlas captures 448,935 cells from 191 high-quality samples across 140 donors and multiple phenotypic groups, identifying 13 distinct cell populations.
- Available at www.pankbase.org, the atlas provides a scalable, rigorously curated platform to support hypothesis-driven diabetes research and can enable a broad range of downstream computational applications.

## Introduction

Type 1 diabetes (T1D) is a complex autoimmune disease characterized by the destruction of insulin-producing beta cells in the pancreatic islets (1,2). Pancreatic islets are composed of multiple specialized endocrine cell types (alpha, beta, delta, gamma, and epsilon cells) that secrete glucose-regulating hormones and dynamically interact with their surrounding microenvironment, including endothelial and immune cells, to maintain blood glucose homeostasis (3).

Single-cell RNA sequencing (scRNA-seq) enables transcriptome profiling at the resolution of individual cells and has become a key approach for studying the cellular mechanisms that contribute to the onset and progression of diabetes (4). The availability of human islets from organ donation coordinated by multiple islet isolation resources, coupled with commercial kits for single-cell sequencing, has enabled numerous scRNA-seq studies of human islets (Supplementary Table 1). However, datasets from individual studies are often modest in size. For example, most 10X Chromium islet scRNA-seq studies published in the past three years were limited to fewer than 20 donors, with considerable overlap in donor and dataset composition across studies, and only two exceptions exceeded 60 donors (5,6) (Supplementary Figure 1, Supplementary Table 1). Although a recent study (7) constructed a large reference map of hormone-secreting cells encompassing a substantial number of cells and donors, its pancreatic tissue data comprised only 18 donors across four studies, underscoring the ongoing challenge of limited donor representation. In addition, inconsistencies in metadata annotation, experimental protocols, and computational workflows across laboratories and data sources hinder meaningful cross-study comparisons and restrict the generalizability of findings. Together, these challenges highlight the critical need for uniform aggregation and harmonization of scRNA-seq datasets from multiple sources to fully realize the scientific potential of available data.

Over the past decade, several resources have expanded access to high-quality islet tissue and associated data. The Integrated Islet Distribution Program (IIDP) has been distributing human islet preparations from multiple isolation centers, using standardized protocols, to investigators worldwide, supporting reproducible islet research (8). The Human Pancreas Analysis Program (HPAP) (9) isolates and profiles pancreatic islets, and shares the resulting datasets through the open-access PancDB resource. In addition, Prodo Labs has been providing isolated human islets to investigators in academia and the pharmaceutical industry to support diabetes research. Together, these resources have enabled the generation of rich datasets that are well positioned for large-scale integration into a community reference.

Here, we present an integrated map of the human pancreatic islet transcriptome, developed by collating publicly available scRNA-seq data generated from islets provided by resources including HPAP, IIDP, and Prodo Labs. Starting from raw sequencing reads rather than aggregating pre-computed count matrices, we applied systematic quality control (QC) of datasets and performed computational integration of high-quality samples to produce a uniformly harmonized reference. We provide this resource through interactive applications that enable researchers to visualize, query, and analyze the atlas, along with open-source pipelines used to generate the reference map and annotations, at www.pankbase.org.

## Research Design and Methods

### Data collection

For the Human Pancreas Analysis Program (HPAP), raw sequencing data and metadata generated from isolated islets using the 10X Chromium platform were downloaded from PancDB at https://hpap.pmacs.upenn.edu/ (accessed March 2024) (6,9). To identify scRNA-seq studies utilizing isolated islets from the Integrated Islet Distribution Program (IIDP) and [Prodo Labs (Prodo)](https://prodolabs.com/), two major US resources for human islet research with well-annotated metadata, we conducted comprehensive literature searches through the Gene Expression Omnibus (GEO; searched before July 2024) (see Supplementary Methods).

### Sample and barcode-level quality control

To confirm donor metadata and identify potential sample swaps, we compared sequencing read profiles with genome-wide genotyping data when available. When genotyping data were unavailable, we performed manual matching using donor age, sex, body mass index (BMI), and allocation center information (see Supplementary Methods).

To systematically assess library quality, we ranked barcodes by decreasing UMI count and identified the change points (“knees”) in each barcode rank profile using a deterministic algorithm based on the Savitzky–Golay filter (10). We retained all libraries with at least two knees. Libraries with a single inferred knee were manually inspected against additional QC metrics before deciding whether to retain them (see Supplementary Methods).

To identify high-quality cell barcodes, we applied a multi-step filtering workflow that included separating cell-containing barcodes from empty droplets, correcting for ambient RNA contamination (11), and filtering based on the fraction of mitochondrial reads (see Supplementary Methods). Samples with more than 200 high-quality cells were retained for downstream analyses.

Doublet detection was performed iteratively to remove doublets and doublet-enriched clusters (see Supplementary Methods).

### Data integration and cell annotation

All data integration was performed using Seurat (12) (v4.4.0). We integrated samples with Harmony (13), correcting for sex, BMI, age, studies, treatments, chemistry and islet distribution resources, iteratively removed low-quality and doublet-enriched clusters, and annotated the remaining clusters to 13 major cell types using canonical markers of pancreatic cell types (see Supplementary Methods).

### Differential expression analysis

Because the map aggregates data from multiple islet distribution resources, isolation centers and studies, we estimated latent factors of technical variation with RUVSeq (14) (v1.36.0) and tested for differentially expressed genes (DEG) using DESeq2 (15) (v1.42.1), accounting for key variables such as sex, age, BMI, ethnicity, chemistry and isolation centers (see Supplementary Methods). We applied a 5% false discovery rate (FDR) to identify differentially expressed genes between T1D and control samples for each cell type.

### Data and Resource Availability

All intermediate analysis results are available at https://data.pankbase.org/analysis-sets/PKBDS1349YHGQ/. The final annotated dataset is available at https://zenodo.org/records/15596314. Interactive visualization of the single-cell map can be accessed at https://pankbase.org/single-cell.html?datasetId=islet_of_Langerhans_scRNA_v3-4.

Intermediate results for differential expression analysis are available at https://data.pankbase.org/browse/ under “Principal Analysis Results”. Final differential expression analysis results can be viewed at https://pankbase.org/diff-exp.html?comparison=dea_comp_1.

Code for scRNA-seq data preprocessing is available at https://github.com/PanKbase/snRNAseq-NextFlow and https://github.com/PanKbase/Multiome-Doublet-Detection-NextFlow. Code for quality control and integration can be found at https://github.com/PanKbase/PanKbase-scRNA-seq. Code for differential expression analysis is available at https://github.com/PanKbase/PanKbase-DEG-analysis.

## Results

We curated 200 scRNA-seq samples from isolated human pancreatic islets acquired through three islet distribution resources: HPAP, IIDP and Prodo Labs (Figure 1A). The transcriptomic data were generated either directly by HPAP or by independent studies conducted on islets distributed by IIDP or Prodo Labs (see Methods for data collection process). To ensure data integrity, we applied a rigorous quality-control (QC) pipeline beginning from raw sequencing reads and assessing quality at the read, cell, and sample levels (see Methods and Supplementary Methods for details).

**Figure 1:**
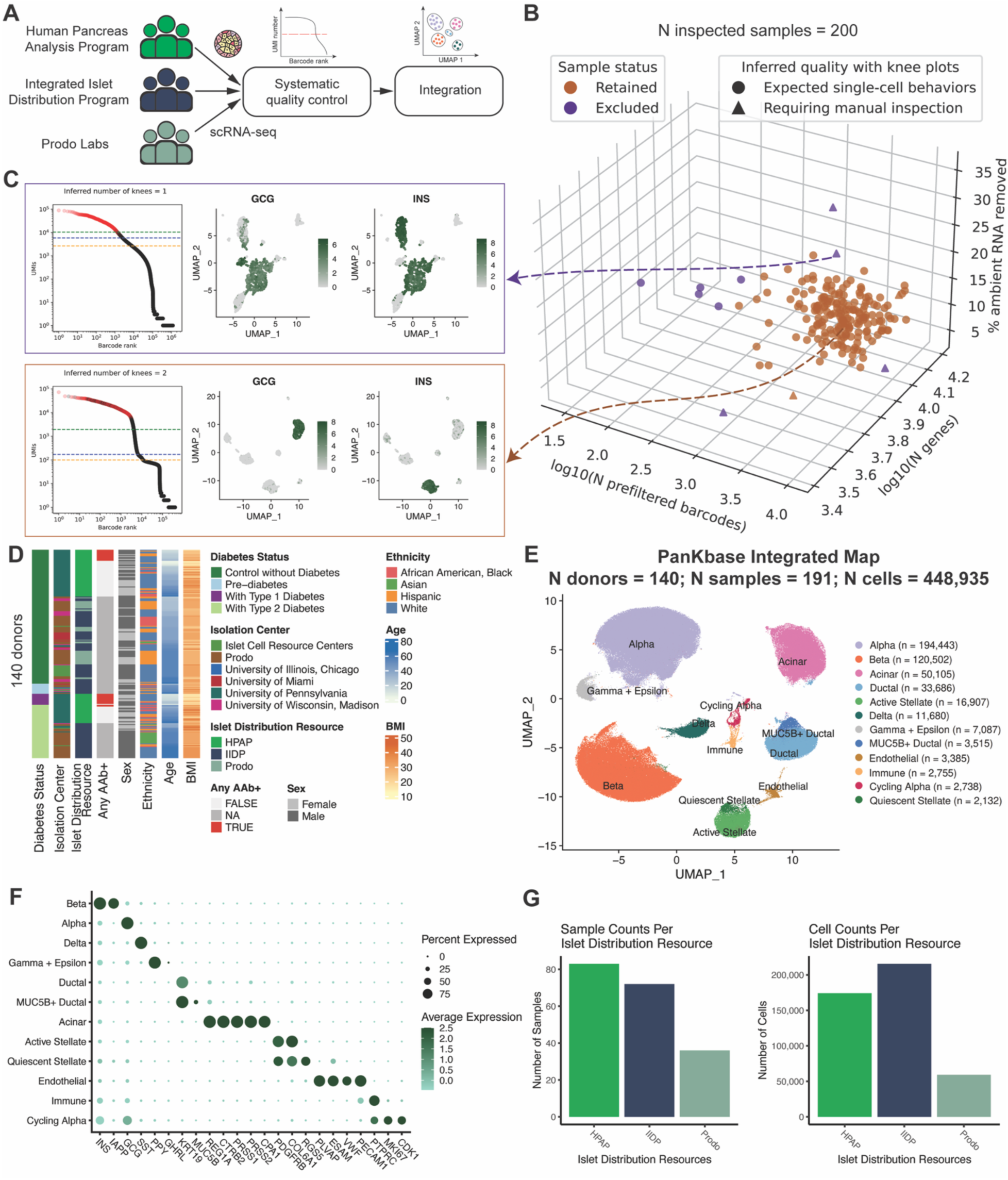
Overview of the PanKbase integrated scRNA-seq atlas of human isolated islets. (A) Study overview. Raw scRNA-seq data generated and/or provided by the Human Pancreas Analysis Program (HPAP), the Integrated Islet Distribution Program (IIDP), and Prodo Labs were aggregated and processed through systematic quality control (QC) and harmonization. (B) Scatterplot summarizing selected QC metrics for the 200 samples evaluated prior to integration. “N prefiltered barcodes” denotes the number of barcodes that passed the per-sample barcode-level EmptyDrops, CellBender, and mitochondrial read-fraction filters (see Supplementary Methods). “% ambient RNA removed” is the maximum fraction of ambient RNA removed during correction (CellBender runs with default settings). “N genes” indicates the maximum number of genes per sample with ≥5 UMIs (computed using prefiltered barcodes). Samples are shaped based on library quality as diagnosed by knee-plot analysis and colored by whether they were retained for downstream analysis. (C) Representative barcode rank plots and marker expression. In each barcode rank plot, each point is a barcode ranked by decreasing UMI count (x-axis: log(rank); y-axis: log(UMIs per barcode)). Barcodes passing barcode-level QC are highlighted (red: pass; black: fail). Green dashed line: inflection point; blue dashed line: end-off-cliff point; yellow dashed line: plateau point. Feature plots show expression levels of INS and GCG among prefiltered singlets (ambient corrected using CellBender with default settings). Libraries inferred to have a single “knee” show poor separation of endocrine marker expression across barcodes, consistent with compromised single-cell quality. (D) Donor metadata. Heatmap summarizing metadata for the 140 donors included in the PanKbase integrated map. “Any AAb+” indicates positivity for at least one autoantibody (GADA, IAA, IA-2, or ZNT8). Abbreviations: HPAP, Human Pancreas Analysis Program; IIDP, Integrated Islet Distribution Program; Prodo, Prodo Labs. (E) Integrated cell atlas. UMAP visualization of 448,935 cells clustered and annotated into 13 cell populations, using data from 140 donors (191 samples). (F) Canonical markers. Dot plot showing normalized expression and the fraction of expressing cells for selected canonical marker genes across clusters. (G) Dataset composition. Bar plots showing the number of samples and cells contributed by each islet distribution resource (HPAP, IIDP, Prodo).

At the sample level, we evaluated each library using barcode rank plots. High-quality libraries typically show two clear change points (“knees”) with a sharp drop between them, reflecting a clean separation between cell-containing droplets and empty droplets. In contrast, low-quality libraries exhibit a blurred transition between these barcode populations, consistent with poor single-cell resolution. To evaluate this diagnostic systematically, we developed a procedure (see Supplementary Methods) that deterministically quantifies the number of knees in each barcode rank plot and flags libraries with a single inferred knee for manual review prior to downstream processing. We then combined knee-based flags with additional QC metrics, including ambient RNA estimates, the estimated number of barcodes and detected genes, and preliminary clustering profiles, to filter libraries and retain samples based on overall quality (Figure 1B; see Methods and Supplementary Methods). Of 193 samples exhibiting expected single-cell behavior, 188 were retained for having ≥200 prefiltered barcodes. Seven samples were flagged for manual inspection; three of these were retained because they demonstrated lower ambient RNA contamination, higher gene detection rates, and well-resolved cell type clusters compared to the four that were excluded (Figure 1B, C). Overall, this QC strategy improves reproducibility and supports scalable library-level assessment.

Ultimately, we assembled an integrated transcriptome of 191 high-quality scRNA-seq samples representing 140 unique donors (59 female, 81 male), creating, to our knowledge, the most comprehensive single-cell atlas of human pancreatic islets to date. The atlas includes 69 donors without diabetes, 12 autoantibody-positive without diabetes, 11 individuals with pre-diabetes, 12 with type 1 diabetes (T1D), and 36 with type 2 diabetes (T2D) (Figure 1D), and includes experimentally perturbed samples, such as SARS-CoV-2 infection and pro-inflammatory cytokine exposure (Supplementary Figure 2). In total, the atlas comprises 448,935 cells which are mapped to 13 populations annotated based on canonical marker genes of pancreatic cell types (Figure 1E, F). Alpha and beta cells represent the largest fractions (43.3% and 26.8%, respectively), while rarer populations in the pancreas, including immune cells (0.6%), are also captured (Figure 1E). We also obtained non-endocrine clusters such as acinar and ductal cells (11.7% and 7.5%, respectively), likely attributable to variability in islet preparation purity. At both the sample and cell levels, the atlas includes contributions from all three islet distribution resources (Figure 1A, G). By integrating across sources, we captured two to five times more samples and cells than any single islet distribution resource alone, expanding the breadth of donors, samples, and cells represented in a unified reference.

As a use case, we performed cell type-specific differential expression analyses comparing donors with T1D versus controls without diabetes. Because the atlas aggregates transcriptomes from multiple studies, it also captures technical heterogeneity that can confound disease-associated signals. For example, all T1D islets in our collection were isolated at the University of Pennsylvania (Figure 1D), whereas islets from donors of other phenotypic groups were obtained from multiple sites, making it challenging to disentangle disease effects (control vs. T1D) from site-specific differences introduced during isolation and processing across islet isolation centers. To improve robustness, we used a latent variable framework (14) to model and adjust for unwanted technical variation prior to differential expression testing (Figure 2A). Briefly, we estimated latent factors of unwanted variation using cell type-specific negative control genes (see Methods and Supplementary Methods), and then examined correlations between these latent variables and donor-, sample-, and cell-level metadata and QC metrics to support interpretation and guide selection of the number of factors (Figure 2A, C). We applied this framework to the six most abundant cell populations: alpha, beta, acinar, ductal, active stellate, and delta cells (Figure 2B). In beta cells, for instance, latent variables were strongly associated with islet isolation center information but showed little association with diabetes status (Figure 2C). We therefore included these latent variables as covariates in our differential expression models to reduce technical confounding while preserving the biological contrast of interest.

**Figure 2:**
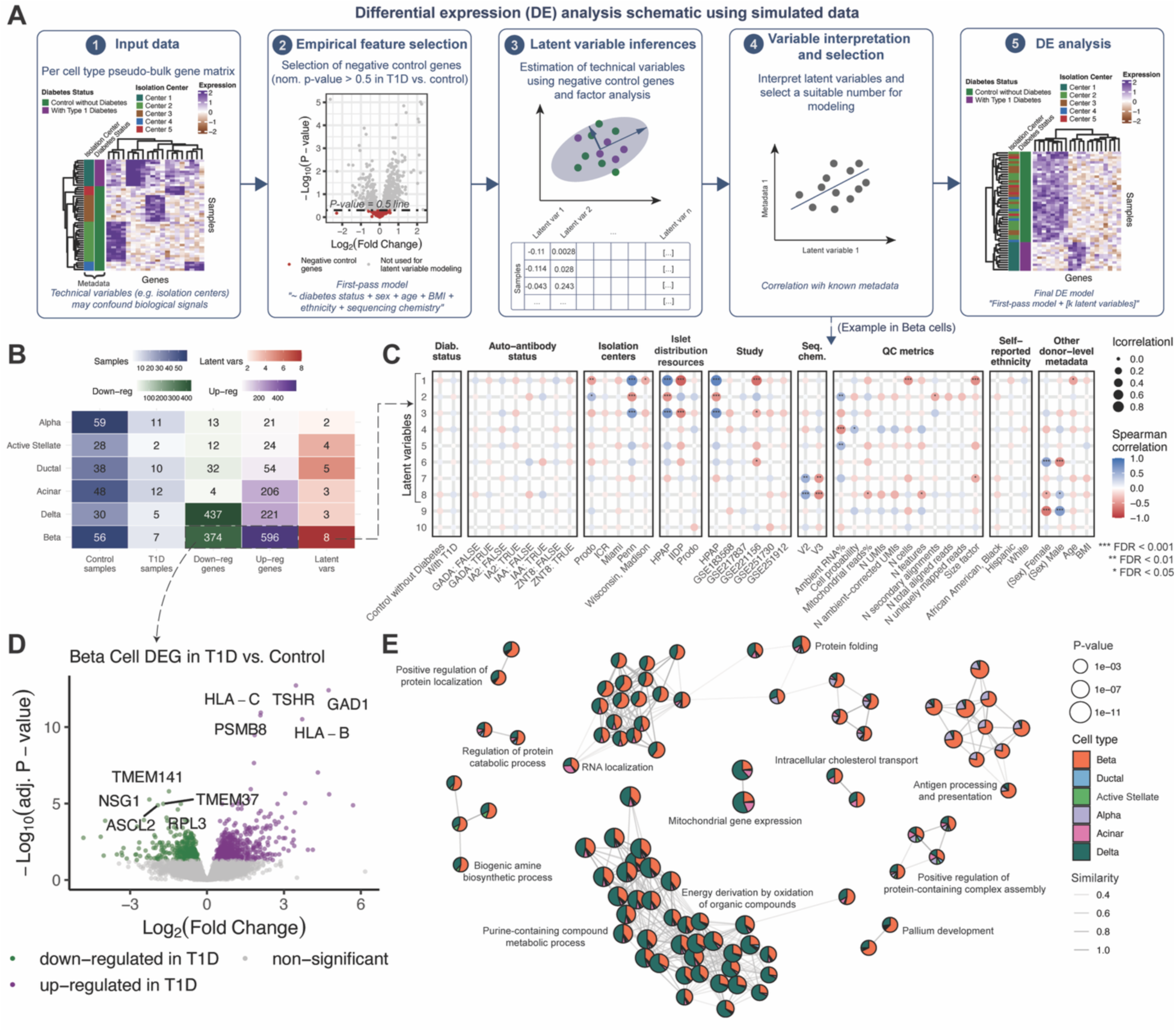
Differential expression analysis comparing Type 1 Diabetes (T1D) vs. control samples. (A) Schematic of a latent variable workflow for differential expression analysis. Observed data can be confounded by strong technical variation such as from isolation centers as T1D samples are confined to a single center. To remove these technical effects, we implemented a latent variable modeling approach. After correction, samples no longer cluster by technical effects, improving the ability to detect underlying biological signals. Diagrams are based on simulated data. Abbreviations: DE, differential expression; vars, variables. (B) Heatmap summarizing, for each cell population, the number of samples, the number of differentially expressed genes, and the number of latent variables included in the model. Abbreviations: up-reg, upregulated; down-reg, downregulated; vars, variables; T1D, type 1 diabetes. (C) Dot plot showing correlations between latent variables and known metadata and QC metrics. In the beta-cell analysis, *“k_stop”* was set to 10 because latent variable 10 showed no correlation with any recorded metadata or QC metric (see Supplementary Methods). We included eight latent variables in the final model, as latent variable 9 did not correlate with any metadata or QC metrics beyond those already captured by the other latent variables. Abbreviations: diab. status, diabetes status; ICR, Islet Cell Resource Center; Miami, University of Miami Sylvester Comprehensive Cancer Center; Penn, University of Pennsylvania; Wisconsin, University of Wisconsin, Madison; BMI, body mass index; QC, quality control; seq. chem., sequencing chemistry. Dot size indicates the absolute correlation magnitude; color indicates the correlation value. For auto-antibody status, correlations were calculated using only samples with available information. (D) Differential expression results in beta cells for T1D vs. control. Upregulated genes are shown in green, downregulated genes in purple, and non-significant genes in grey (FDR ≥ 5%). The top five most significant genes in both directions are labeled. (E) Gene Ontology enrichment network showing a subset of enriched terms from the DEG sets. Edges represent term similarity, and node colors indicate the percent of observed genes from each cell type. Terms with the highest number of observed genes are labeled.

Across the six cell types analyzed, we identified 1,805 unique genes that were differentially expressed between controls and individuals with T1D at FDR < 5% (Supplementary Table 2). Beta cells showed the largest transcriptional change, with 596 genes up-regulated and 374 down-regulated in T1D (970 total; Figure 2B, D; Supplementary Table 2). Prominent up-regulated genes included MHC class I genes (e.g., *HLA-B*, *HLA-C*), *GAD1*, and *TSHR*, consistent with prior reports (5,16). In comparison, fewer differentially expressed genes (DEGs) were detected in other populations (Figure 2B; Supplementary Table 2). We benchmarked these DEG sets against those reported in Fasolino et al. (16) and Elgamal et al. (5), two recent studies that performed the same T1D vs. control comparisons. Using the same FDR threshold (5%) and pseudobulk approach, the PanKbase analysis identified more DEGs and generally larger effect-size estimates, while remaining highly concordant with both studies in the direction of effects (Supplementary Figures 3 and 4). This pattern is consistent with increased statistical power arising from the larger number of donors contributing to the pseudobulk profiles.

Pathway enrichment of these DEGs implicated protein homeostasis, mitochondrial function, and immune recognition, with antigen processing and presentation terms driven largely by alpha and beta cells (Figure 2E; Supplementary Table 3). Cell-type-specific immune pathways, including leukocyte-mediated immunity and T cell-mediated cytotoxicity, were up-regulated in T1D in alpha and beta cells (Supplementary Figures 5 and 6; Supplementary Table 3), indicating pronounced immune-related transcriptional changes in these cell types in T1D.

## Discussion

In this study, we present an integrated single-cell atlas of isolated human islets which comprises 191 samples from 140 donors across three major human islet research resources. The atlas contains major endocrine cell types such as alpha and beta cells, and non-endocrine populations, e.g. ductal and acinar cells, which are retained to varying degrees during the purification process. Raw data were obtained from publicly available resources and processed through rigorous, systematic QC and harmonization, yielding a comprehensive, cell type-resolved view of transcriptomic variation within islet tissue. The atlas supports hypothesis-driven discovery in islet biology and diabetes research, and serves as a reference for cell-type annotation (17) and integrated analyses in future studies. We placed particular emphasis on dataset curation and QC to ensure quality consistency and completeness. These properties are essential for model development and training (18,19), including foundation-model construction (20) and other AI-enabled analyses.

Moreover, we processed all datasets starting from raw sequencing reads and applied a uniform QC workflow, rather than relying on study-specific filtered matrices, reducing the risk of introducing study-dependent biases during preprocessing. Specifically, aggregating pre-filtered matrices inherits each study’s choices of reference, cell calling, ambient-RNA correction and doublet removal, which cannot be reversed downstream and leave technical variation confounded with biology at integration (13,21,22). Because ambient contamination varies across experiments and produces off-target expression of one cell type’s markers in another, inconsistently processed matrices can yield spurious multi-hormonal or doublet-driven populations that persist through batch correction, which uniform reprocessing from raw reads can help remove (11,21,23). Importantly, the QC procedures described here are tissue-agnostic and are publicly released for broader use in single-cell and single-nucleus RNA-seq studies.

This study has certain limitations worth noting. First, despite the large cell count, gamma and epsilon cells remain clustered together. Although epsilon cells are detectable by scRNA-seq in principle (24), their rarity (25) and the technical heterogeneity inherent in multi-source data likely are still major challenges that current computational tools cannot yet reliably resolve.

Second, while the atlas greatly expands coverage of control samples and T2D donor samples across all distribution resources, samples from donors with T1D remain limited to HPAP. Incorporating additional T1D (4) or T1D-simulating datasets (26) in future iterations will help address this imbalance. Furthermore, complementary data modalities such as functional and imaging data available through the Network for Pancreatic Organ Donors with Diabetes (nPOD) (27), the Alberta Diabetes Institute IsletCore (ADI) (28) and through cross-cutting resources such as Pancreastlas (29) and others (30), presents an exciting opportunity for subsequent versions. Meanwhile, the expanded representation of control and T2D samples enables more detailed characterization of these groups, providing a clear direction for future work to understand changes in T2D.

A key methodological contribution is a latent variable-based analytical framework that applies established methods (14) to account for unwanted technical variation in a multi-source atlas. Here, we developed an interpretable pipeline to select and examine the latent variables based on their correlations with known metadata and QC metrics, enabling more transparent adjustment for potential confounders. We identified many genes with altered expression in T1D, highlighting pathways related to protein homeostasis/metabolism and immune-mediated processes. We further benchmarked these results against Fasolino et al. (16) and Elgamal et al. (5) which performed a similar comparison. Overall, we observed concordance in the direction of effects, while this study identified a larger number of significant genes at the pseudobulk level. A subset of genes was detected only in the previous studies, which may reflect differences between studies in QC filters, ambient RNA correction methods, gene filtering and differential testing strategies.

In summary, this comprehensive single-cell atlas of isolated human pancreatic islets, supported by carefully curated metadata and rigorous QC, serves as a valuable reference for hypothesis-driven studies and machine-learning applications in islet biology and diabetes research.

## Acknowledgements

We sincerely thank all organ donors and their families. This work used data acquired from the database (https://hpap.pmacs.upenn.edu/) of the Human Pancreas Analysis Program (HPAP; RRID:SCR_016202; PMID: 31127054; PMID: 36206763; PMID: 35228745 and PMID: 37188822). HPAP is part of a Human Islet Research Network (RRID:SCR_014393) consortium (UC4-DK112217, U01-DK123594, UC4-DK112232, and U01-DK123716). Additional islet samples and tissues were provided by the Integrated Islet Distribution Program (IIDP) (RRID:SCR_014387), funded by NIH Grant #2UC4DK098085.

During the course of preparing this work, the author(s) used Claude for the purpose of polishing some sections of the writing. Following the use of this tool/service, the author(s) formally reviewed the content for its accuracy and edited it as necessary. The author(s) take full responsibility for all the content of this publication.

## Author Contributions

Data collection and curation: HTHV, FF, KJG, MB, ALG, SCJP; Metadata collection: HTHV, PK, YS, KJG; Data quality control: HTHV, HS, SS, KJG, ALG, SCJP; Data integration: HTHV, SCJP; Differential expression analysis: HTHV, SCJP, LB, KJG, TB; Writing, original draft: HTHV, SCJP; Review and discussion: all; Supervision: SCJP, KJG, ALG. SCJP is the guarantor of this work and, as such, had full access to all the data in the study and takes responsibility for the integrity of the data and the accuracy of the data analysis.

## References

1. Syed FZ. Type 1 Diabetes Mellitus. Ann Intern Med. 2022 Mar 15;175(3):ITC33–48. doi:10.7326/AITC202203150

2. Chiang JL, Kirkman MS, Laffel LMB, Peters AL, on behalf of the Type 1 Diabetes Sourcebook Authors. Type 1 Diabetes Through the Life Span: A Position Statement of the American Diabetes Association. Diabetes Care. 2014 Jun 12;37(7):2034–54. doi:10.2337/dc14-1140

3. Weitz J, Menegaz D, Caicedo A. Deciphering the Complex Communication Networks That Orchestrate Pancreatic Islet Function. Diabetes. 2020 Dec 15;70(1):17–26. doi:10.2337/dbi19-0033

4. Melton R, Jimenez S, Elison W, Tucciarone L, Howell A, Wang G, et al. Single-cell multiome and spatial profiling reveals pancreas cell type–specific gene regulatory programs of type 1 diabetes progression. Sci Adv. 2025 Sep 10;11(37):eady0080. doi:10.1126/sciadv.ady0080

5. Elgamal RM, Kudtarkar P, Melton RL, Mummey HM, Benaglio P, Okino ML, et al. An Integrated Map of Cell Type–Specific Gene Expression in Pancreatic Islets. Diabetes. 2023 Aug 15;72(11):1719–28. doi:10.2337/db23-0130

6. Patil AR, Schug J, Naji A, Kaestner KH, Faryabi RB, Vahedi G. Single-cell expression profiling of islets generated by the Human Pancreas Analysis Program. Nat Metab. 2023 May 15;5(5):713–5. doi:10.1038/s42255-023-00806-x

7. Fei L, Huang-Doran I, Lawler K, Yu Y, Pett JP, Méndez-Acevedo KM, et al. A Hormone Cell Atlas maps the human endocrine system at cellular resolution. Science. 2026 May 28;0(0):eaeb2672. doi:10.1126/science.aeb2672

8. Brissova M, Niland JC, Cravens J, Olack B, Sowinski J, Evans-Molina C. The Integrated Islet Distribution Program answers the call for improved human islet phenotyping and reporting of human islet characteristics in research articles. Diabetologia. 2019 Jul 1;62(7):1312–4. doi:10.1007/s00125-019-4876-3

9. Kaestner KH, Powers AC, Naji A, HPAP Consortium, Atkinson MA. NIH Initiative to Improve Understanding of the Pancreas, Islet, and Autoimmunity in Type 1 Diabetes: The Human Pancreas Analysis Program (HPAP). Diabetes. 2019 Jul 1;68(7):1394–402. doi:10.2337/db19-0058

10. Savitzky Abraham, Golay MJE. Smoothing and Differentiation of Data by Simplified Least Squares Procedures. Anal Chem. 1964 Jul 1;36(8):1627–39. doi:10.1021/ac60214a047

11. Fleming SJ, Chaffin MD, Arduini A, Akkad AD, Banks E, Marioni JC, et al. Unsupervised removal of systematic background noise from droplet-based single-cell experiments using CellBender. Nat Methods. 2023 Sep;20(9):1323–35. doi:10.1038/s41592-023-01943-7

12. Hao Y, Stuart T, Kowalski MH, Choudhary S, Hoffman P, Hartman A, et al. Dictionary learning for integrative, multimodal and scalable single-cell analysis. Nat Biotechnol. 2024 Feb;42(2):293–304. doi:10.1038/s41587-023-01767-y

13. Korsunsky I, Millard N, Fan J, Slowikowski K, Zhang F, Wei K, et al. Fast, sensitive and accurate integration of single-cell data with Harmony. Nat Methods. 2019 Dec;16(12):1289–96. doi:10.1038/s41592-019-0619-0

14. Risso D, Ngai J, Speed TP, Dudoit S. Normalization of RNA-seq data using factor analysis of control genes or samples. Nat Biotechnol. 2014 Sep;32(9):896–902. doi:10.1038/nbt.2931

15. Love MI, Huber W, Anders S. Moderated estimation of fold change and dispersion for RNA-seq data with DESeq2. Genome Biol. 2014 Dec 5;15(12):550. doi:10.1186/s13059-014-0550-8

16. Fasolino M, Schwartz GW, Patil AR, Mongia A, Golson ML, Wang YJ, et al. Single-cell multi-omics analysis of human pancreatic islets reveals novel cellular states in type 1 diabetes. Nat Metab. 2022 Feb 28;4(2):284–99. doi:10.1038/s42255-022-00531-x

17. Aran D, Looney AP, Liu L, Wu E, Fong V, Hsu A, et al. Reference-based analysis of lung single-cell sequencing reveals a transitional profibrotic macrophage. Nat Immunol. 2019 Feb;20(2):163–72. doi:10.1038/s41590-018-0276-y

18. Subramanyam A, Chen Y, Grossman RL. Scaling Laws Revisited: Modeling the Role of Data Quality in Language Model Pretraining [Internet]. arXiv; 2026 [cited 2026 Apr 16]. Available from: http://arxiv.org/abs/2510.03313 doi:10.48550/arXiv.2510.03313

19. Mohammed S, Budach L, Feuerpfeil M, Ihde N, Nathansen A, Noack N, et al. The effects of data quality on machine learning performance on tabular data. Inf Syst. 2025 Jul 1;132:102549. doi:10.1016/j.is.2025.102549

20. Hao M, Gong J, Zeng X, Liu C, Guo Y, Cheng X, et al. Large-scale foundation model on single-cell transcriptomics. Nat Methods. 2024 Aug;21(8):1481–91. doi:10.1038/s41592-024-02305-7

21. Janssen P, Kliesmete Z, Vieth B, Adiconis X, Simmons S, Marshall J, et al. The effect of background noise and its removal on the analysis of single-cell expression data. Genome Biol. 2023 Jun 19;24(1):140. doi:10.1186/s13059-023-02978-x

22. Lun ATL, Riesenfeld S, Andrews T, Dao TP, Gomes T, Participants in the 1st Human Cell Atlas Jamboree, et al. EmptyDrops: distinguishing cells from empty droplets in droplet-based single-cell RNA sequencing data. Genome Biol. 2019 Dec;20(1):63. doi:10.1186/s13059-019-1662-y

23. Wolock SL, Lopez R, Klein AM. Scrublet: Computational Identification of Cell Doublets in Single-Cell Transcriptomic Data. Cell Syst. 2019 Apr 24;8(4):281–291.e9. doi:10.1016/j.cels.2018.11.005 PubMed PMID: 30954476.

24. Bandesh K, Motakis E, Nargund S, Kursawe R, Selvam V, Ansarullah, et al. Deep single-cell decoding of human pancreatic islets reveals T2D β-cell gene expression defects. EMBO J. 2026 Jun 1;45(11):3978–4005. doi:10.1038/s44318-026-00744-w

25. Andralojc KM, Mercalli A, Nowak KW, Albarello L, Calcagno R, Luzi L, et al. Ghrelin-producing epsilon cells in the developing and adult human pancreas. Diabetologia. 2009 Mar 1;52(3):486–93. doi:10.1007/s00125-008-1238-y

26. Albanus RD, Zhang X, Zhao Z, Taylor HJ, Tang X, Han Y, et al. Integrative single-cell multi-omics profiling of human pancreatic islets identifies T1D-associated genes and regulatory signals. Cell Rep. 2025 Aug 26;44(8). doi:10.1016/j.celrep.2025.116065 PubMed PMID: 40737125.

27. Campbell-Thompson M, Wasserfall C, Kaddis J, Albanese-O’Neill A, Staeva T, Nierras C, et al. Network for Pancreatic Organ Donors with Diabetes (nPOD): Developing a Tissue Biobank for Type 1 Diabetes. Diabetes Metab Res Rev. 2012 Oct;28(7):608–17. doi:10.1002/dmrr.2316 PubMed PMID: 22585677; PubMed Central PMCID: PMC3456997.

28. Lyon J, Manning Fox JE, Spigelman AF, Kim R, Smith N, O’Gorman D, et al. Research-Focused Isolation of Human Islets From Donors With and Without Diabetes at the Alberta Diabetes Institute IsletCore. Endocrinology. 2016 Feb 1;157(2):560–9. doi:10.1210/en.2015-1562

29. Saunders DC, Messmer J, Kusmartseva I, Beery ML, Yang M, Atkinson MA, et al. Pancreatlas: Applying an Adaptable Framework to Map the Human Pancreas in Health and Disease. Patterns. 2020 Oct 5;1(8):100120. doi:10.1016/j.patter.2020.100120 PubMed PMID: 33294866; PubMed Central PMCID: PMC7691395.

30. Chen SY, Lekka C, Lemvig J, Yi X, Cnop M, Marchetti P, et al. From pancreas and islet resources to diabetes insights. Diabetologia. 2026 Jul 1;69(7):1741–58. doi:10.1007/s00125-026-06731-4

