## Supplementary material for "PanKbase Integrated Single-Cell Map: A Comprehensive Atlas of Human Pancreatic Islets": Online Supplementary Materials

#### Table of Content

Supplementary Methods
- Data collection (continued)
- Read-level data processing and quality control
- Sample-level metadata quality control (continued)
- Sample and barcode-level quality control (continued)
- Doublet detection
- Data integration and cell annotation
- Differential expression analysis (continued)
- Pathway enrichment analysis

Supplementary Figures

Supplementary References

### Supplementary Methods

#### Data collection (continued)

Literature searches were conducted to identify original studies involving scRNA-seq generated using islets distributed by IIDP and Prodo. Briefly, pancreatic islet datasets of expression profiling by high throughput sequencing were identified via GEO search for the date range before July 2024, then studies with scRNA-seq data were determined manually. Raw sequencing data was obtained from the following sources: https://hpap.pmacs.upenn.edu/ (1–4), GSE142465 (5), GSE159556 (6), GSE183568 (7), GSE190447 (8), GSE201256 (9), GSE217837 (10), GSE221156 (11), GSE251730 (12), GSE251912 (13). Raw data outside of PancDB were downloaded using the *fastq-dump* command from the SRA toolkit (14) (v3.1.1).

To identify studies utilizing 10X Chromium scRNA-seq data from human pancreas or isolated islets (see Supplementary Table 1, Supplementary Figure 1), we conducted a literature search using the keywords 'scRNA-seq' and 'islets', restricted to publications from 2023 through April 2026, with sample sizes for each study determined through manual curation.

#### Read-level data processing and quality control

The quality of raw reads was assessed using FastQC (15) (v0.11.9), followed by aggregation of individual sample reports with MultiQC (16) (v1.11). Comprehensive inspection of these reports enabled the identification of technical artifacts, including flow cell irregularities, base calling errors, and unexpected technical sequence contamination. Sequencing runs were included for further analysis if more than 50% of tiles, as indicated by the Per Tile Quality and Quality Score plots, demonstrated high quality (i.e., quality scores at or above the average across all bases).

Alignment of raw reads to the human genome (hg38) was performed using STARsolo (17) (v2.7.10a), employing comprehensive gene annotation from Gencode release 39 (https://www.gencodegenes.org/human/release_39.html) to generate STAR index files (STAR --*runMode genomeGenerate*, default parameters). This process yielded count matrices quantifying features such as gene expression levels. For downstream analysis, GeneFull_ExonOverIntron count matrices were utilized. MultiQC was also employed to summarize STARsolo log files, facilitating assessment of key sequencing metrics (e.g., percentage of uniquely mapped reads, overall mapping rates). High-quality reads were extracted using SAMtools (18) (v1.14) with the following command: *samtools view -h -b -q 255 -F 4 -F 256 -F 2048 $bam.*

To process hashtag-oligoes samples (HTO), HTO matrix files and barcode-treatment maps were downloaded from the GEO page ID: GSE251912 (13). Then, the HTODemux() function from the Seurat package (19) (v4.4.0) was used to demultiplex the cells to their original treatments.

#### Sample-level metadata quality control (continued)

To confirm donor metadata and check for sample swaps, we compared the read profiles with genotyping data using the mbv tool (20) included in QTLtools (21) (v1.3.1). When donor unique identifiers were not supplied in the original studies, we assigned donor identities to samples when *perc_het_consistent* and *perc_hom_consistent* from mbv are both greater than 0.8. When donors did not have genotype data, we manually searched for individuals with matching age, sex, BMI and islet isolation centers to assign donor identifiers to samples. Subsequently, we retained only samples with a determined identifier. Donor phenotypic groups were defined in accordance with the classifications reported in the respective origin studies (1–10,12,13,22).

#### Sample and barcode-level quality control (continued)

To systematically inspect quality metrics underlying all libraries, we utilized the Unique Molecular Identifiers (UMI) numbers and their ranks using the following algorithm. We ranked every barcode $b$ based on their UMI numbers ($n_b$) in a decreasing order, then considered $log(n_b)$ as a function of the rank's logarithm. Next, we calculated the first-order discrete difference between two consecutive barcodes to obtain the rate of changes in $n_b$ along the rank axis, then smoothed out this difference signal profile using the Savitzky–Golay filter (23). As a result, we identified points along the profile with the largest change rates, which aligned with the “knees”, and quantified the “knee” point UMI numbers.

To distinguish potential cells from empty droplets, EmptyDrops (24) from the DropletUtils (25) (v1.26.0) was used. Droplets with significant deviations from the ambient profile at FDR < 0.005 were regarded as non-empty.

Utilizing the mathematical framework underlying EmptyDrops, we identified the "end-of-cliff point" on barcode rank plots for each sample. The end-of-cliff point is characterized as the position with a log-rank value surpassing all detected inflection points (as determined by EmptyDrops) and yielding the minimum signed curvature, and was used for subsequent ambient RNA correction.

Ambient RNA contamination was addressed using CellBender (26) (v0.3.0) in two iterative rounds. The first round was run with default settings where correction was performed with a false positive rate (FPR) of 0.05, while all other parameters were set at their default values. The second round was run with modified parameters as follows: --expected_cells was set to the number of estimated true cells, and --total_droplets_included was set to the average number of UMIs observed between the inflection point and the end-of-cliff point.

In order to identify barcodes that correspond to cells, we defined the following criteria: (1) probability that a barcode corresponds to a true cell (obtained from CellBender using default settings) $\geq 0.99$; (2) cells were detected as non-empty using EmptyDrops; (3) cells with fractions of ambient reads < a dynamic threshold determined per sample using the Multi-Otsu Thresholding algorithm on non-empty cells detected using EmptyDrops; and (4) cells with percent of mitochondria chromosome (chrMT) reads < a dynamic threshold determined per sample using the Multi-Otsu Thresholding algorithm on the combination of two sets: non-empty cells, and cells whose chrMT read fraction < 0.3 and rank higher than that of end-of-cliff points.

#### Doublet detection

For each sample, DoubletFinder (27) (v2.0.3) was used twice on the RNA ambient-corrected gene count matrix obtained using modified runs of CellBender, and the second round of DoubletFinder was run without doublets detected in the first round. Next, we jointly clustered libraries across each tissue source (HPAP, IIDP and Prodo) using Seurat with doublets included, using Harmony (28) (v1.2.0) to correct for the following covariates: sex, BMI, age, origin studies, treatments and sequencing chemistry, and resolution of 1.8. We removed doublets and doublet-enriched clusters which are defined as ones with doublet rates > 65%.

#### Data integration and cell annotation

All data integration was performed using Seurat (v4.4.0). In particular, due to the large number of samples, we first combined all samples per islet distribution resources (data generated using HPAP, IIDP, and Prodo samples) to create three Seurat objects, retaining cells that expressed at least one protein-coding gene, and protein-coding genes that are expressed in at least one cell. Subsequently, we merged all objects and carried out integration using Harmony, correcting for the following covariates: sex, BMI, age, origin studies, treatments, sequencing chemistry, and islet distribution resources. As a result, we obtained a set of 481,154 cells expressing 19,546 protein-coding genes.

Next, we adopted an iterative approach to further clean up the data. Firstly, we clustered the cells (resolution = 1.8), and tested for clusters with significantly different profiles of gene numbers and UMI numbers using Wilcoxon rank sum test. Here, we define a fold change as median(tested cluster's metrics) / median(all other clusters' metrics). A cluster is determined to be significantly different if their adjusted p-value < 0.05 and fold change > 2, and they are removed from the subsequent analysis. As a result, we obtained a set of 476,429 cells, expressing 19,546 protein-coding genes. Secondly, we focused on identifying doublet-like cells using the remaining cells. For each sample, we began by clustering cells at a resolution of 4. We pinpointed clusters that were enriched with cells exhibiting UMI counts higher than expected and expressing markers from at least two cell populations in 50% or more of the cells. To achieve this, we performed Binomial tests to assess whether a cluster had more cells with UMI counts exceeding the 90th percentile of the samples than the expected rate of 0.1, using a Benjamini-Hochberg adjusted P-value threshold of < 0.05. Cells within these enriched clusters were labeled as "doublet-like cells." Next, we employed Fisher's exact test to identify clusters in the integrated map that were enriched with doublet-like cells (Benjamini-Hochberg adjusted P-value threshold of < 0.05). Clusters in the integrated map that both were enriched with doublet-like cells and expressed markers from at least two cell populations in 50% or more of the cells were subsequently removed. Additionally, all doublet-like cells were removed. Third, we re-clustered and harmonized the data (resolution = 1.8), resulting in 48 clusters. Finally, using the following list of markers, we annotated the cell clusters to 13 major cell types: INS and IAPP (Beta cells); GCG (Alpha cells); SST (Delta cells); PPY (Gamma cells); GHRL (Epsilon cells); CFTR and KRT19 (Ductal cells); REG1A, CTRB2, PRSS1, PRSS2, CPA1 (Acinar cells); PDGFRB, COL6A1 (Activated stellate cells); PDGFRB, COL6A1, and RGS5 (Quiescent stellate cells); PECAM1, PLVAP, ESAM, VWF (Endothelial cells); PTPRC (Immune cells); GCG, MKI67, CDK1 (Cycling alpha cells).

#### Differential expression analysis (continued)

As the PanKbase single cell map consists of a wide collection of data from multiple tissue sources, isolation centers and independent studies, we used a latent variable analysis to discern biological variation from technical variation as we conducted analyses of differentially expressed genes (DEG). Using DESeq2 (29) (v1.42.1), we compared gene expression profiles between type

1 diabetes (T1D) and control samples for every cell type with more than 20 samples and for each sample with more than 20 non-treated cells; one sample per donor was randomly selected when replicates existed. To minimize potential effects of known and unknown confounding factors, we included known covariates in the DESeq2 model as well as accounted for unknown covariates using the RUVSeq latent variable approach (30) (v1.36.0). In brief, we used the following multi-step process. First, we removed genes from the raw count matrix that had less than 10 reads in fewer than 25% of the samples for that cell type. Second, we ran a first-pass differential expression analysis using DESeq2 with sex, age, BMI, ethnicity and sequencing chemistry as known covariates. The output result was filtered for genes that were not differentially expressed between T1D and control samples which had nominal p values > 0.5. These genes were used as negative control genes for the RUVSeq::RUVg function to estimate latent variables accounting for variation in the data not attributed to disease status. Third, the latent variables estimated from the RUVseq run were then used as additional covariates (in addition to sex, age, BMI, ethnicity and chemistry) for the second run of DESeq2.

To select the number of latent variables $k$, we applied the following algorithm: (1) we carried out latent variable estimation for a wide range of variable numbers (total tested $k \leq 30$); (2) out of all tested variables, we define a $k$ named "*k_stop*", which is a $k$ that either significantly correlates with diabetes status or is the smallest $k$ that does not significantly correlate with any known variables (Spearman correlation, Benjamini-Hochberg adj. p value < 0.05); (3) from $k = 1$ to $k = k\_stop$, we selected the smallest $k$ such that it correlates with at least one unique known variable that is not identified by any other $k$.

We applied a 5% false discovery rate (FDR) to identify differentially expressed genes between T1D and control samples for each cell type.

#### Pathway enrichment analysis

To determine the relevant functions of the DEGs across all six analyzed cell types, we used gene ontology (GO) analysis on the combined gene lists. clusterProfiler (31) (v4.10.1) was used, with the human annotation from the org.Hs.eg.db R package (32) (v3.18.0). Similarity across GO terms was assessed using the function pairwise_termsim() from enrichplot R package (33) (v1.22.0). A GO term was considered enriched when its adjusted p-value ≤ 0.05.

On a cell-type level, we carried out gene set enrichment analysis to resolve directional changes of pathways using clusterProfiler (31) (v4.10.1) functions gseKEGG() and gseGO(). For this analysis, all genes (sorted based on their log2(fold change)) were used. A pathway was considered enriched when its adjusted p-value ≤ 0.05.

### Supplementary Figures

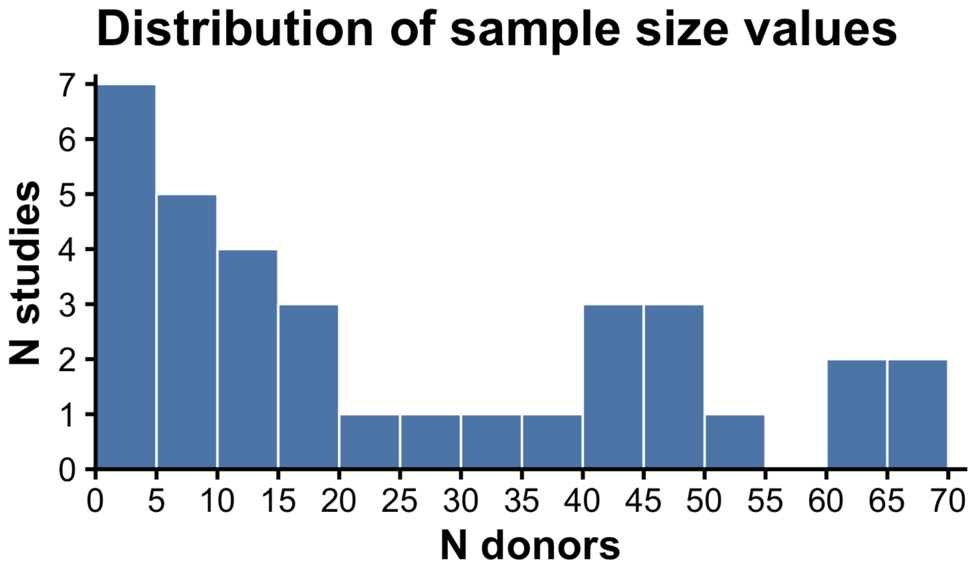


Supplementary Figure 1: Distribution of sample sizes across studies with 10X Chromium scRNA-seq data from human pancreas or isolated islets. See Supplementary Table 1 for the full list of publications.

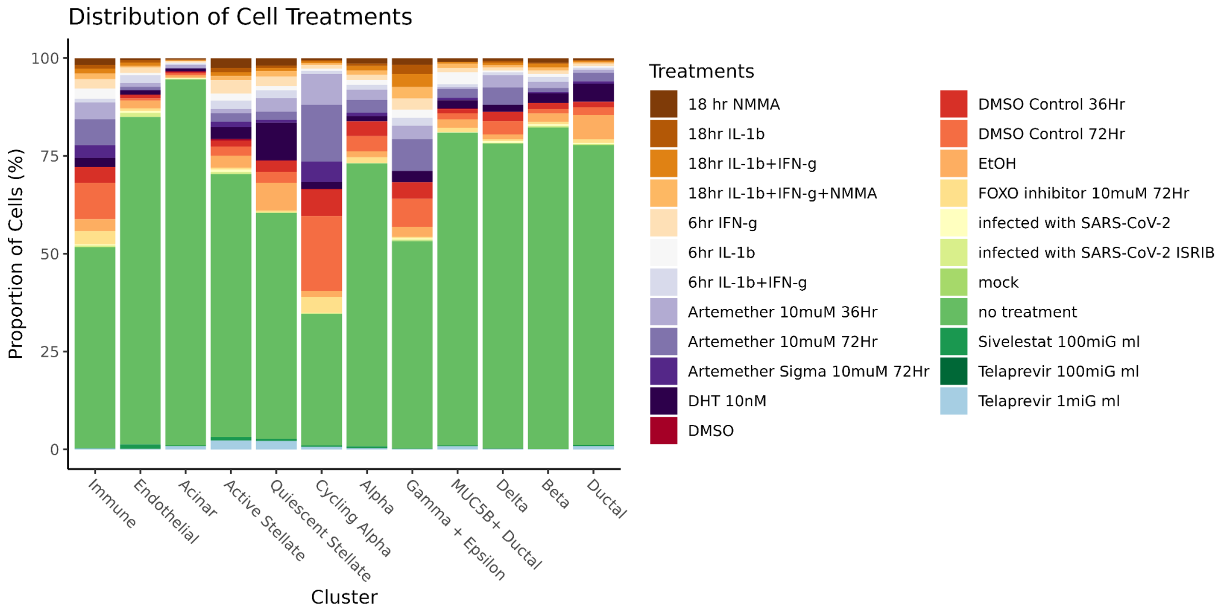


Supplementary Figure 2: Proportion of cells in each treatment condition, stratified by cell population.

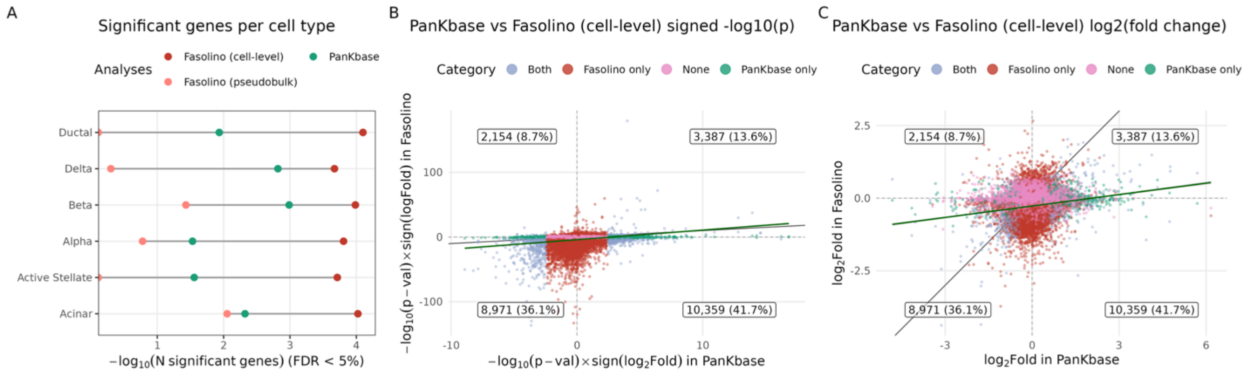


Supplementary Figure 3: Benchmarking differential expression results from the PanKbase integrated map against Fasolino et al.

(A) Bubble plot showing the total number of significantly differentially expressed genes (FDR < 5%) from Fasolino et al. (cell- and pseudobulk-level analyses) and the PanKbase analysis (pseudobulk-level only). At the pseudobulk level, the PanKbase analysis identifies more significant genes, whereas at the cell level, the Fasolino et al. analysis identifies more.

(B) Scatter plot comparing results for beta cells across genes tested in both studies using signed -log10(nominal p values). Each point represents a gene and is colored by significance category (FDR < 5%): significant in both studies (green), Fasolino-only (orange), PanKbase-only (pink), or not significant in either (purple). The best-fit regression line is shown in solid green and the identity line in solid black. Overall, genes show strong concordance in the direction of effects across studies.

(C) Scatter plot comparing beta-cell effect sizes across genes tested in both studies using log2(fold change). Points are colored as in (B); the best-fit regression line is shown in solid green and the identity line in solid black. The regression indicates larger estimated effect sizes in the PanKbase analysis for the same genes.

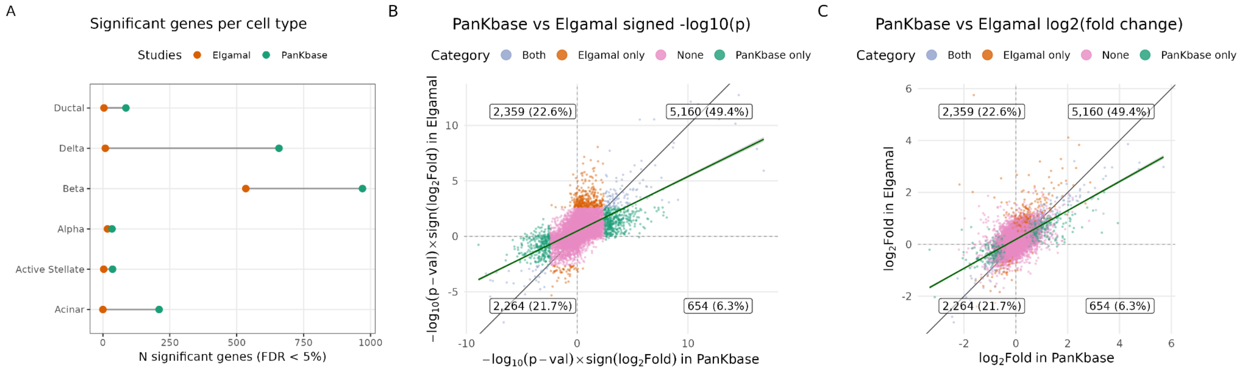


Supplementary Figure 4: Benchmarking differential expression results from the PanKbase integrated map against Elgamal et al.

(A) Bubble plot showing the total number of significantly differentially expressed genes (FDR < 5%) identified in Elgamal et al. and in the PanKbase analysis. The PanKbase-based analysis identifies more genes.

(B) Scatter plot comparing results for beta cells across genes tested in both studies using signed -log10(nominal p values). Each point represents a gene and is colored by significance category (FDR < 5%): significant in both studies (green), Elgamal-only (orange), PanKbase-only (pink), or not significant in either (purple). The best-fit regression line is shown in solid green and the identity line in solid black. Overall, genes show strong concordance in the direction of effects across studies.

(C) Scatter plot comparing beta-cell effect sizes across genes tested in both studies using log2(fold change). Points are colored as in (B); the best-fit regression line is shown in solid green and the identity line in solid black. The regression indicates larger estimated effect sizes in the PanKbase analysis for the same genes.

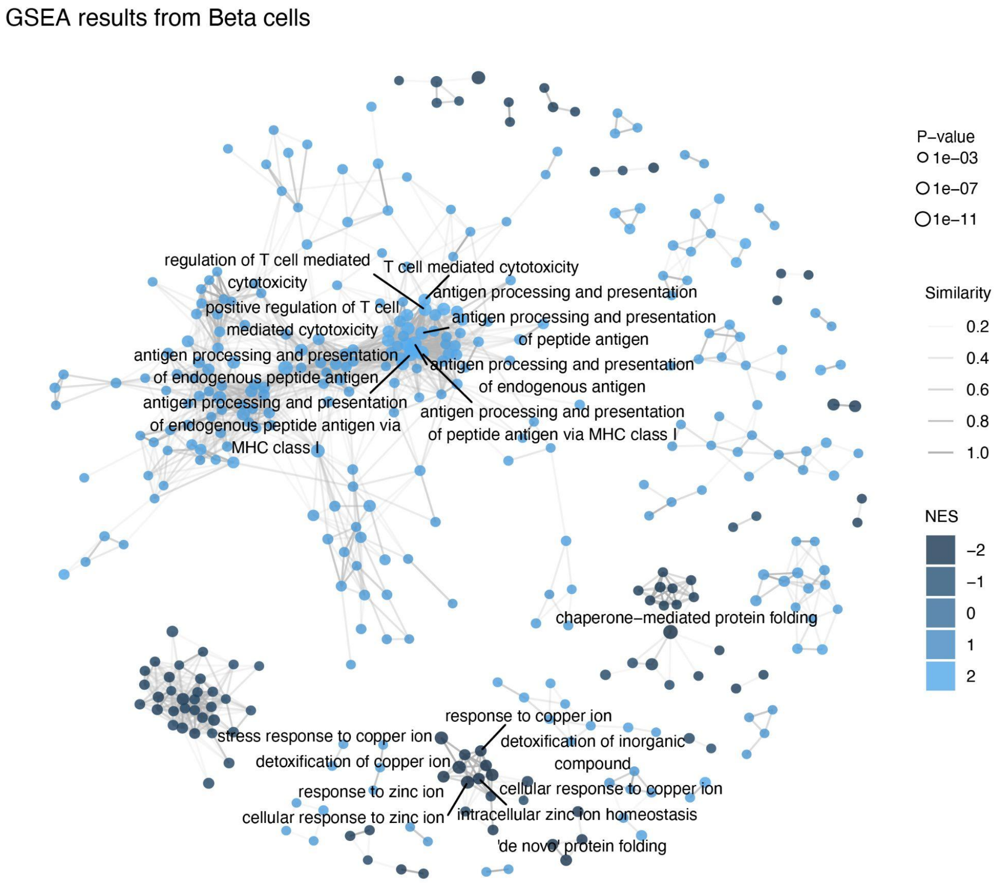


Supplementary Figure 5: Gene set enrichment analysis (GSEA) network showing a subset of enriched terms in beta cells. Edges represent term similarity, and node colors indicate normalized enrichment scores (NES). The top 10 positively and negatively enriched terms are labeled.

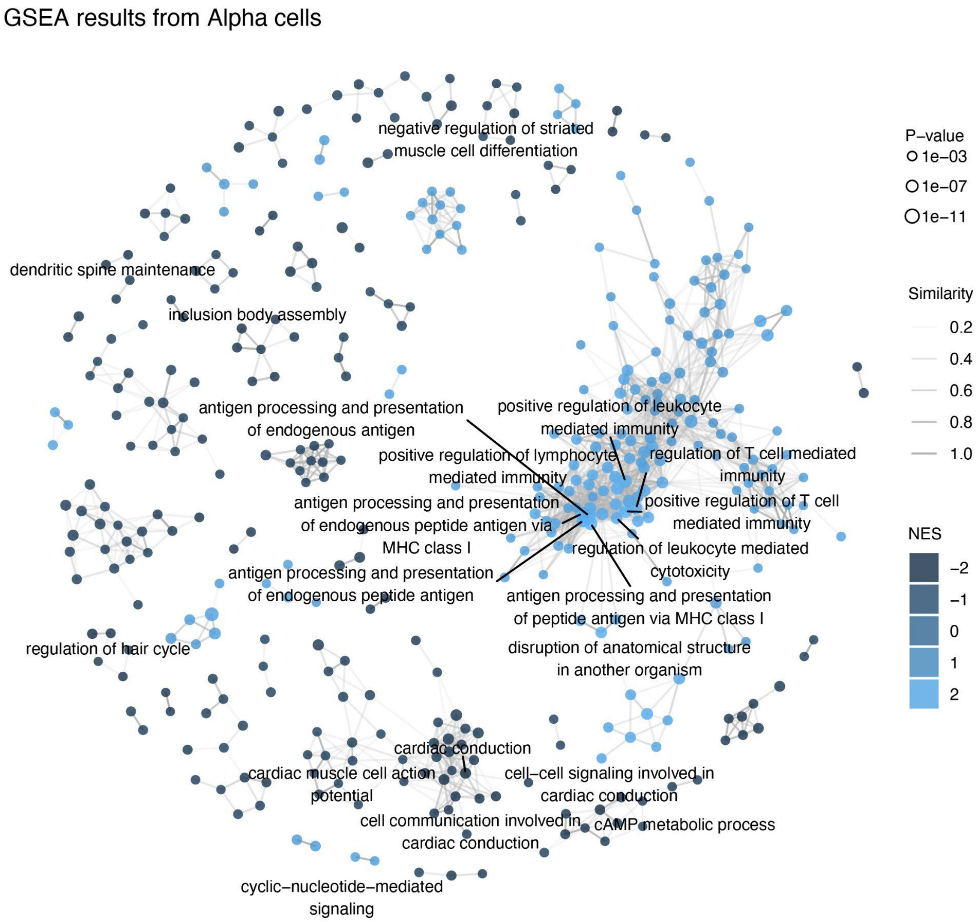


Supplementary Figure 6: Gene set enrichment analysis (GSEA) network showing a subset of enriched terms in alpha cells. Edges represent term similarity, and node colors indicate normalized enrichment scores (NES). The top 10 positively and negatively enriched terms are labeled.

### Supplementary References


1. Kaestner KH, Powers AC, Naji A, HPAP Consortium, Atkinson MA. NIH Initiative to Improve Understanding of the Pancreas, Islet, and Autoimmunity in Type 1 Diabetes: The Human Pancreas Analysis Program (HPAP). Diabetes. 2019 Jul 1;68(7):1394–402. doi:10.2337/db19-0058

2. Patil AR, Schug J, Naji A, Kaestner KH, Faryabi RB, Vahedi G. Single-cell expression profiling of islets generated by the Human Pancreas Analysis Program. Nat Metab. 2023 May 15;5(5):713–5. doi:10.1038/s42255-023-00806-x

3. Fasolino M, Schwartz GW, Patil AR, Mongia A, Golson ML, Wang YJ, et al. Single-cell multi-omics analysis of human pancreatic islets reveals novel cellular states in type 1 diabetes. Nat Metab. 2022 Feb 28;4(2):284–99. doi:10.1038/s42255-022-00531-x

4. Shapira SN, Naji A, Atkinson MA, Powers AC, Kaestner KH. Understanding islet dysfunction in type 2 diabetes through multidimensional pancreatic phenotyping: The Human Pancreas Analysis Program. Cell Metab. 2022 Dec;34(12):1906–13. doi:10.1016/j.cmet.2022.09.013

5. Marquina-Sanchez B, Fortelny N, Farlik M, Vieira A, Collombat P, Bock C, et al. Single-cell RNA-seq with spike-in cells enables accurate quantification of cell-specific drug effects in pancreatic islets. Genome Biol. 2020 Dec;21(1):106. doi:10.1186/s13059-020-02006-2

6. Tang X, Uhl S, Zhang T, Xue D, Li B, Vandana JJ, et al. SARS-CoV-2 infection induces beta cell transdifferentiation. Cell Metab. 2021 Aug;33(8):1577-1591.e7. doi:10.1016/j.cmet.2021.05.015

7. Shrestha S, Saunders DC, Walker JT, Camunas-Soler J, Dai XQ, Haliyur R, et al. Combinatorial transcription factor profiles predict mature and functional human islet α and β cells. JCI Insight. 2021 Sep 22;6(18):e151621. doi:10.1172/jci.insight.151621

8. Basile G, Vetere A, Hu J, Ijaduola O, Zhang Y, Liu KC, et al. Excess pancreatic elastase alters acinar-β cell communication by impairing the mechano-signaling and the PAR2 pathways. Cell Metab. 2023 Jul;35(7):1242-1260.e9. doi:10.1016/j.cmet.2023.05.007

9. Xu W, Qadir MMF, Nasteska D, Mota De Sa P, Gorvin CM, Blandino-Rosano M, et al. Architecture of androgen receptor pathways amplifying glucagon-like peptide-1 insulinotropic action in male pancreatic β cells. Cell Rep. 2023 May;42(5):112529. doi:10.1016/j.celrep.2023.112529

10. Kang RB, Li Y, Rosselot C, Zhang T, Siddiq M, Rajbhandari P, et al. Single-nucleus RNA sequencing of human pancreatic islets identifies novel gene sets and distinguishes β-cell subpopulations with dynamic transcriptome profiles. Genome Med. 2023 May 1;15(1):30. doi:10.1186/s13073-023-01179-2

11. Bandesh K, Motakis E, Nargund S, Kursawe R, Selvam V, Ansarullah, et al. Deep single-cell decoding of human pancreatic islets reveals T2D β-cell gene expression defects. EMBO J. 2026 Jun 1;45(11):3978–4005. doi:10.1038/s44318-026-00744-w

12. Stancill JS, Kasmani MY, Cui W, Corbett JA. Single Cell RNAseq Analysis of Cytokine-Treated Human Islets: Association of Cellular Stress with Impaired Cytokine Responsiveness. Function. 2024 Jul 11;5(4):zqae015. doi:10.1093/function/zqae015

13. Sokolowski EK, Kursawe R, Selvam V, Bhuiyan RM, Thibodeau A, Zhao C, et al. Multi-omic human pancreatic islet endoplasmic reticulum and cytokine stress response mapping provides type 2 diabetes genetic insights. Cell Metab. 2024 Nov;36(11):2468-2488.e7. doi:10.1016/j.cmet.2024.09.006

14. SRA Toolkit Development Team. SRA Toolkit [Internet]. Available from: https://trace.ncbi.nlm.nih.gov/Traces/sra/sra.cgi?view=software

15. Andrews S. FastQC [Internet]. Available from: https://www.bioinformatics.babraham.ac.uk/projects/fastqc/

16. Ewels P, Magnusson M, Lundin S, Käller M. MultiQC: summarize analysis results for multiple tools and samples in a single report. Bioinformatics. 2016 Oct 1;32(19):3047–8. doi:10.1093/bioinformatics/btw354

17. Kaminow B, Yunusov D, Dobin A. STARsolo: accurate, fast and versatile mapping/quantification of single-cell and single-nucleus RNA-seq data [Internet]. Bioinformatics; 2021 [cited 2025 May 30]. Available from: http://biorxiv.org/lookup/doi/10.1101/2021.05.05.442755 doi:10.1101/2021.05.05.442755

18. Li H, Handsaker B, Wysoker A, Fennell T, Ruan J, Homer N, et al. The Sequence Alignment/Map format and SAMtools. Bioinformatics. 2009 Aug 15;25(16):2078–9. doi:10.1093/bioinformatics/btp352

19. Hao Y, Hao S, Andersen-Nissen E, Mauck WM, Zheng S, Butler A, et al. Integrated analysis of multimodal single-cell data. Cell. 2021 Jun;184(13):3573-3587.e29. doi:10.1016/j.cell.2021.04.048

20. Fort A, Panousis NI, Garieri M, Antonarakis SE, Lappalainen T, Dermitzakis ET, et al. *MBV*: a method to solve sample mislabeling and detect technical bias in large combined genotype and sequencing assay datasets. Stegle O, editor. Bioinformatics. 2017 Jun 15;33(12):1895–7. doi:10.1093/bioinformatics/btx074

21. Delaneau O, Ongen H, Brown AA, Fort A, Panousis NI, Dermitzakis ET. A complete tool set for molecular QTL discovery and analysis. Nat Commun. 2017 May 18;8(1):15452. doi:10.1038/ncomms15452

22. Bandesh K, Motakis E, Nargund S, Kursawe R, Selvam V, Bhuiyan RM, et al. Single-cell decoding of human islet cell type-specific alterations in type 2 diabetes reveals converging genetic- and state-driven β-cell gene expression defects [Internet]. Genomics; 2025 [cited 2025 May 30]. Available from: http://biorxiv.org/lookup/doi/10.1101/2025.01.17.633590 doi:10.1101/2025.01.17.633590

23. Savitzky Abraham, Golay MJE. Smoothing and Differentiation of Data by Simplified Least Squares Procedures. Anal Chem. 1964 Jul 1;36(8):1627–39. doi:10.1021/ac60214a047

24. Lun ATL, Riesenfeld S, Andrews T, Dao TP, Gomes T, Participants in the 1st Human Cell Atlas Jamboree, et al. EmptyDrops: distinguishing cells from empty droplets in droplet-based single-cell RNA sequencing data. Genome Biol. 2019 Dec;20(1):63. doi:10.1186/s13059-019-1662-y

25. Aaron Lun JG. DropletUtils [Internet]. Bioconductor; 2018 [cited 2025 May 30]. Available from: https://bioconductor.org/packages/DropletUtils doi:10.18129/B9.BIOC.DROPLETUTILS

26. Fleming SJ, Chaffin MD, Arduini A, Akkad AD, Banks E, Marioni JC, et al. Unsupervised removal of systematic background noise from droplet-based single-cell experiments using CellBender. Nat Methods. 2023 Sep;20(9):1323–35. doi:10.1038/s41592-023-01943-7

27. McGinnis CS, Murrow LM, Gartner ZJ. DoubletFinder: Doublet Detection in Single-Cell RNA Sequencing Data Using Artificial Nearest Neighbors. Cell Syst. 2019 Apr;8(4):329-337.e4. doi:10.1016/j.cels.2019.03.003

28. Korsunsky I, Millard N, Fan J, Slowikowski K, Zhang F, Wei K, et al. Fast, sensitive and accurate integration of single-cell data with Harmony. Nat Methods. 2019 Dec;16(12):1289–96. doi:10.1038/s41592-019-0619-0

29. Love MI, Huber W, Anders S. Moderated estimation of fold change and dispersion for RNA-seq data with DESeq2. Genome Biol. 2014 Dec 5;15(12):550. doi:10.1186/s13059-014-0550-8

30. Risso D, Ngai J, Speed TP, Dudoit S. Normalization of RNA-seq data using factor analysis of control genes or samples. Nat Biotechnol. 2014 Sep;32(9):896–902. doi:10.1038/nbt.2931

31. Yu G, Wang LG, Han Y, He QY. clusterProfiler: an R Package for Comparing Biological Themes Among Gene Clusters. OMICS J Integr Biol. 2012 May;16(5):284–7. doi:10.1089/omi.2011.0118 PubMed PMID: 22455463; PubMed Central PMCID: PMC3339379.

32. Carlson M. Bioconductor [Internet]. [cited 2026 Apr 8]. org.Hs.eg.db: Genome wide annotation for Human. Available from: http://bioconductor.org/packages/org.Hs.eg.db/ doi:10.18129/B9.bioc.org.Hs.eg.db

33. Bioconductor [Internet]. [cited 2026 Jun 1]. enrichplot. Available from: http://bioconductor.org/packages/enrichplot/
